# Genomic Disaggregation Reveals Distinct Admixture Patterns and Cardiometabolic Risk Loci in Black Hawaiians

**DOI:** 10.64898/2026.01.24.701518

**Authors:** Kasra Vand, Nelson Badía, Bohdan B. Khomtchouk

## Abstract

**Background:** The systematic aggregation of distinct admixed subpopulations into broad racial categories creates genomic blind spots that undermine the promise of precision medicine. Black Hawaiians (BH) exemplify this exclusion. Characterized by a unique tri-continental ancestry (African, European, and Native Hawaiian/Pacific Islander) and disproportionate cardiometabolic burden, their population-specific risk drivers remain masked by systematic conflation with broader ancestral cohorts.

**Methods:** We performed the first comprehensive genomic analysis of 287 BH participants from the NIH *All of Us* Research Program using whole-genome sequencing (WGS). Following haplotype phasing (SHAPEIT5), we characterized population structure (ADMIXTURE, PCA), inferred local ancestry tracts (RFMix), and reconstructed demographic history (SMC++). Genome-wide allele frequency differentiation (AFD) was calculated against tri-continental reference panels, and Electronic Health Record (EHR) data were integrated to quantify the population’s cardiometabolic burden.

**Results:** The cohort exhibited complex tri-continental admixture (mean: 67.0% African, 22.1% European, 10.9% NHPI) with high inter-individual heterogeneity. Phenotypic analysis confirmed a substantial disease burden (34.8% hypertension, mean BMI 31.2 kg/m^2^), while SMC++ reconstruction revealed a sharp demographic bottleneck in recent generations. Genome-wide AFD analysis of 8.9M variants demonstrated systematic differentiation (mean Δ vs African: 0.041, NHPI: 0.069, European: 0.084). The top 100 differentiated variants mapped to 31 unique genes, identifying distinct candidates including *MYO9A*, RAB37, and *PEAR1*. Notably, differentiation in the cytoskeletal regulator *MYO9A* suggests a mechanostructural etiology for kidney disease distinct from classical *APOL1* cytotoxicity, while *PEAR1* variants implicate population-specific pharmacogenomic resistance to antiplatelet therapy.

**Conclusion:** This study highlights the critical necessity of data disaggregation in genomic research, using the Black Hawaiian population as a paradigmatic example. By distinguishing this community from broader aggregate groups, we uncovered a distinct genomic architecture with unique admixture patterns that drive specific cardiometabolic risks. These findings demonstrate the necessity of granular resolution for achieving equitable precision medicine.

## 1. Introduction

The success of precision medicine hinges on accurately characterizing the genetic architecture of disease across all global populations. However, despite increasing investment in regional research initiatives, the majority of globally aggregated genomic data continues to overrepresent individuals from broader, less admixed racial categories, a disparity that is amplified in complexly admixed communities. These populations, formed by the recent interbreeding of two or more distinct ancestral groups, possess unique genetic landscapes that offer both scientific opportunities and ethical obligations for focused research. Black Hawaiians (BH), also known as Pōpolo, represent a historically and genetically distinct population resulting from tri-continental admixture involving African, European, and Native Hawaiian/Pacific Islander (NHPI) ancestries. Despite comprising a recognized demographic group with a unique sociocultural history in Hawai’i, BH individuals have been consistently aggregated with broader and genetically less relevant racial categories, such as African American or general NHPI groups, in large-scale genomic initiatives. This exclusion perpetuates health disparities by preventing the identification of population-specific genetic risk factors and limiting the predictive power of polygenic risk scores (PRS) in this community. This population is of particular significance due to its elevated cardiometabolic disease burden, including disproportionate rates of hypertension (HTN), type 2 diabetes (T2D), and obesity.

Admixed populations provide a powerful framework for admixture mapping, a technique that leverages the distinct linkage disequilibrium patterns created by recent ancestry mixing events to efficiently fine-map disease-associated loci. Realizing this potential, however, is contingent upon the availability of population-specific genomic reference data that accurately characterize local ancestry tracts and allele frequency distributions. Leveraging whole-genome sequencing (WGS) data and comprehensive phenotyping from the NIH *All of Us* Research Program (AoU), this study presents the first systematic and multi-faceted genomic characterization of BH individuals (N=287). Our primary objectives were to: (1) determine global and local ancestry proportions across the tri-continental components (African, European, NHPI) and assess inter-individual heterogeneity; (2) utilize Sequentially Markov Coalescent (SMC++) to reconstruct the recent effective population size (*N*_*e*_) history, providing a temporal context for admixture and population bottlenecks; (3) conduct genome-wide allele frequency differentiation (AFD) analysis relative to ancestral populations to identify variants with unique frequencies that may underlie population-specific cardiometabolic risk.

This study establishes the foundational genomic architecture of BH and identifies systematically differentiated candidate loci. These findings provide critical resources for advancing equitable precision medicine initiatives in this community and serve as a model for disaggregated research in other underrepresented admixed groups.

## 2. Methods

### 2.1. Study Population and Data Acquisition

Black Hawaiian individuals were identified from the NIH AoU database through intersectional querying of self-reported race and ethnicity. Participants who self-identified as both “Black or African American” AND “Native Hawaiian or Other Pacific Islander” were included. Of 413 initially identified individuals, 287 had available whole-genome sequencing data and passed the quality control and were included in the final analysis cohort. Whole-genome sequencing data were generated using Illumina short-read technology with mean coverage >30×. Individual VCF files were accessed through the AoU Researcher Workbench and processed locally for population genetic analyses. Phenotypic data including clinical diagnoses, laboratory measurements, and physical measurements were extracted using the dataset builder within the Researcher Workbench.

### 2.2. Genomic Data Processing and Quality Control

Individual VCF files were merged using bcftools^1^ (v1.12) and converted to PLINK^2^ binary format for quality control. Standard quality control procedures were applied: exclusion of variants with >5% missing data, minor allele frequency <1%, Hardy-Weinberg equilibrium p<1×10^−6^, and individuals with >5% missing genotypes. Following quality control, the dataset comprised 287 individuals (out of 413) and 16.4 million variants with 99.8% genotyping rate. For population structure analyses, variants were pruned for linkage disequilibrium (LD) using a sliding window approach (50-variant windows, 5-variant step, r^2^<0.5), yielding approximately 2.2 million quasi-independent markers. All autosomal chromosomes were phased using SHAPEIT5^3^ with the 1000 Genomes Project (1KGP) Phase 3 reference panel^4^ and GRCh38 genetic maps.

### 2.3. Population Structure Analysis

Principal component analysis was performed using PLINK^2^ (v1.9) on LD-pruned variants to assess population structure. The first 20 principal components were calculated and visualized to evaluate clustering patterns and identify potential stratification. Global ancestry proportions were estimated using ADMIXTURE^5^ (v1.3.0) with K values ranging from 2 to 6. Cross-validation error was used to determine the optimal number of ancestral populations. Results are presented for K=3 and K=5 to capture both major ancestral components and finer population structure.

### 2.4. Local Ancestry Inference

Local ancestry inference was performed using RFMix^6^ (v2.03) to estimate ancestry tract lengths and ancestry-specific allele frequencies along the genome. Reference panels were constructed from 1KGP populations representing African (Yoruba, n=108), European (CEU, n=99), and NHPI ancestry, which we constructed by extracting and phasing genomes of individuals who self-identified exclusively as Native Hawaiian or Other Pacific Islander from the AoU database (n=300), ensuring no overlap with the Black Hawaiian study cohort. All reference panels and BH query genomes were phased using SHAPEIT5. To accommodate computational constraints, variants were thinned to every 5th position, yielding ∼200,000 variants per chromosome while maintaining sufficient marker density for accurate ancestry inference. RFMix was run with default parameters using a three-way admixture model (African, European, NHPI). Genome-wide ancestry proportions were calculated by averaging local ancestry calls across all chromosomes for each individual.

### 2.5. Demographic History Inference

To reconstruct the effective population size (*N*_*e*_) history of the Black Hawaiian population over time, we utilized SMC++^7^, a sequentially Markovian coalescent method that leverages LD patterns in whole-genome sequence data. This method is particularly well-suited for unphased or phased genomes and can handle sample sizes in the hundreds, making it ideal for our cohort (N=287). We performed the analysis on the autosomes using the composite likelihood approach. The mutation rate was set to the standard human rate of μ = 1.25 * 10^-8^ mutations per base pair per generation, and a generation time of 29 years was assumed. The analysis estimated the trajectory of *N*_*e*_ from the present day back to approximately 35,000 generations ago.

### 2.6. Allele Frequency Differentiation Analysis

To identify variants with unusual frequency patterns in BH, we calculated allele frequencies in the study cohort and compared them to reference populations (African/YRI, European/CEU, and NHPI). Allele frequencies were computed using bcftools for all variants present in both the BH dataset and reference panels (8.9 million overlapping variants). For each variant, we calculated the absolute allele frequency difference (Δ) between BH and each reference population. The maximum Δ across all three comparisons was used as the primary measure of differentiation. The top 100 most differentiated variants were annotated for gene location using the UCSC Genome Browser API to identify genes harboring variants with unusual frequency patterns.

### 2.7. Phenotype Data Extraction and Analysis

Clinical phenotype data were extracted from electronic health records within the AoU database. Diagnoses of T2D, cardiovascular disease, and HTN were identified using ICD-10 codes. Laboratory measurements (HDL cholesterol, glucose, HbA1c, triglycerides) and physical measurements (BMI, blood pressure) were extracted where available. Summary statistics were calculated for all phenotypes to characterize the cardiometabolic burden in the study population.

### 2.8. Statistical Analysis

All analyses were performed in Python 3.10 and R 4.2. Descriptive statistics are reported as means ± standard deviations for continuous variables and counts (percentages) for categorical variables. Data visualization was performed using matplotlib and seaborn. All genomic coordinates are reported in GRCh38/hg38.

## 3. Results

### 3.1. Cohort Characteristics and Cardiometabolic Burden

The final study cohort comprised 287 Black Hawaiian individuals with whole-genome sequencing data. Phenotypic characterization revealed a substantial cardiometabolic disease burden. All 278 individuals had available measurement data, and 100 individuals (34.8%) had diagnosed hypertension, 22 (7.7%) had T2D, and 18 (6.3%) had cardiovascular disease. Mean BMI was 31.2 kg/m^2^ (obese category), with mean systolic blood pressure of 129.9 mmHg and diastolic blood pressure of 80.9 mmHg. Laboratory measurements were available for 98-195 individuals depending on the specific biomarker, with mean HDL cholesterol of 51.1 mg/dL, glucose of 104.0 mg/dL, and triglycerides of 126.9 mg/dL (Figure 1).

**Figure 1.**
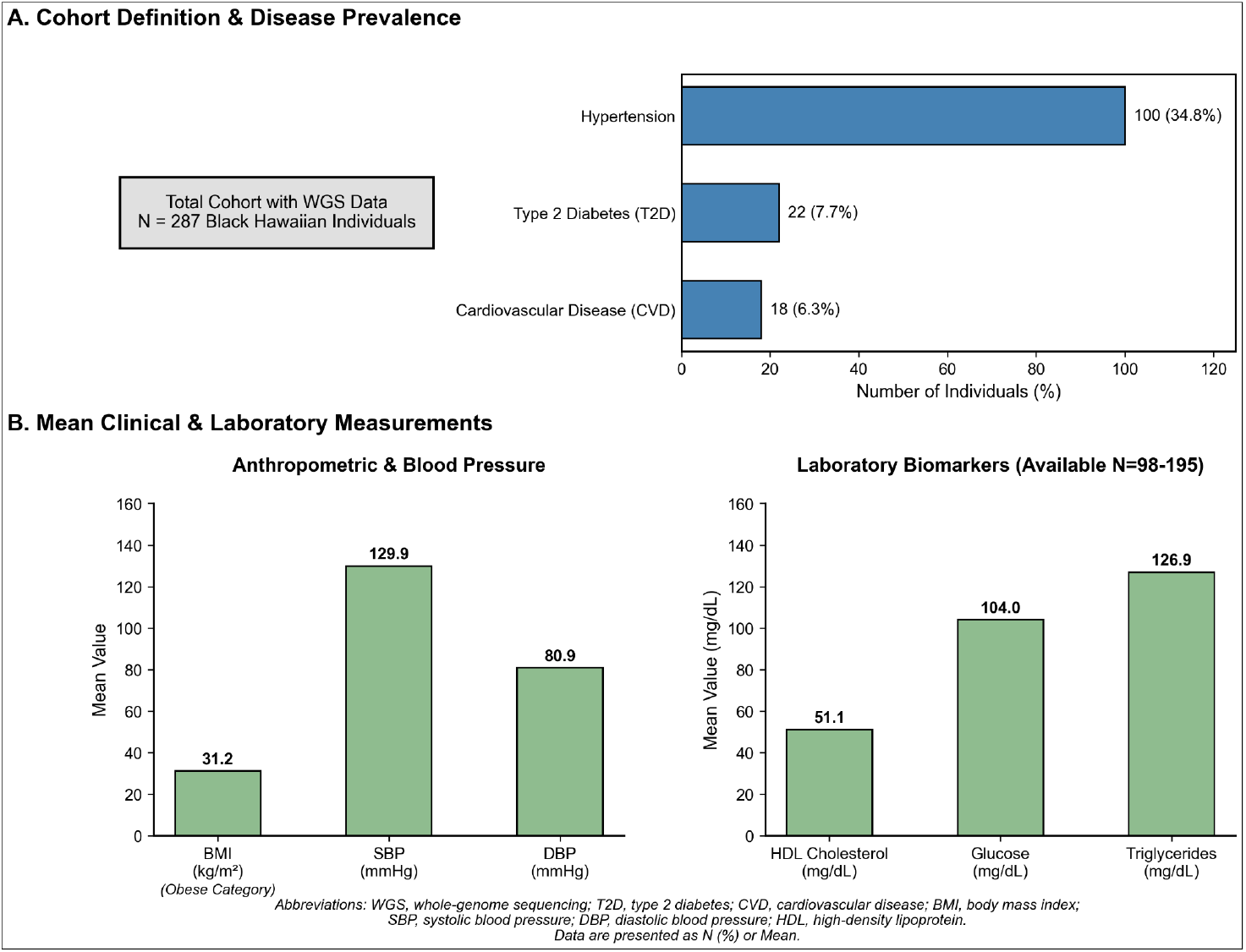
Clinical Characteristics of the Black Hawaiian Study Cohort (N=287). This two-panel figure summarizes the cardiometabolic disease burden of the study participants. Panel A depicts the cohort definition flowchart and presents the prevalence rates for diagnosed hypertension, type 2 diabetes, and cardiovascular disease. Panel B displays mean values for key anthropometric measurements, blood pressure readings, and available laboratory biomarkers, highlighting mean BMI in the obese category.

### 3.2. Complex Tri-Continental Admixture Patterns

Principal component analysis (PCA) revealed that BH individuals occupy a distinct region of genetic space intermediate between African, European, and Pacific Islander reference populations, consistent with tri-continental admixture (Figure 2-A). No discrete sub-clusters were observed within the BH cohort, indicating continuous variation in ancestry proportions rather than distinct ancestral groups. Local ancestry inference using RFMix provided chromosome-specific ancestry assignments, revealing mean genome-wide ancestry proportions of 67.0% African (±26.9%), 22.1% European (±20.9%), and 10.9% NHPI (±19.3%) (as an example Figure 3 shows the local ancestry for chromosome 2). The substantial standard deviations reflect high inter-individual heterogeneity in admixture proportions, with no clear population substructure (Figure 2-B). ADMIXTURE analysis at K=5 revealed five ancestral components with mean proportions of 23.5%, 17.6%, 18.9%, 22.9%, and 17.1% respectively (Figure 2-C). Cross-validation indicated K=5 as optimal, suggesting fine-scale population structure beyond the three major continental groups.

**Figure 2.**
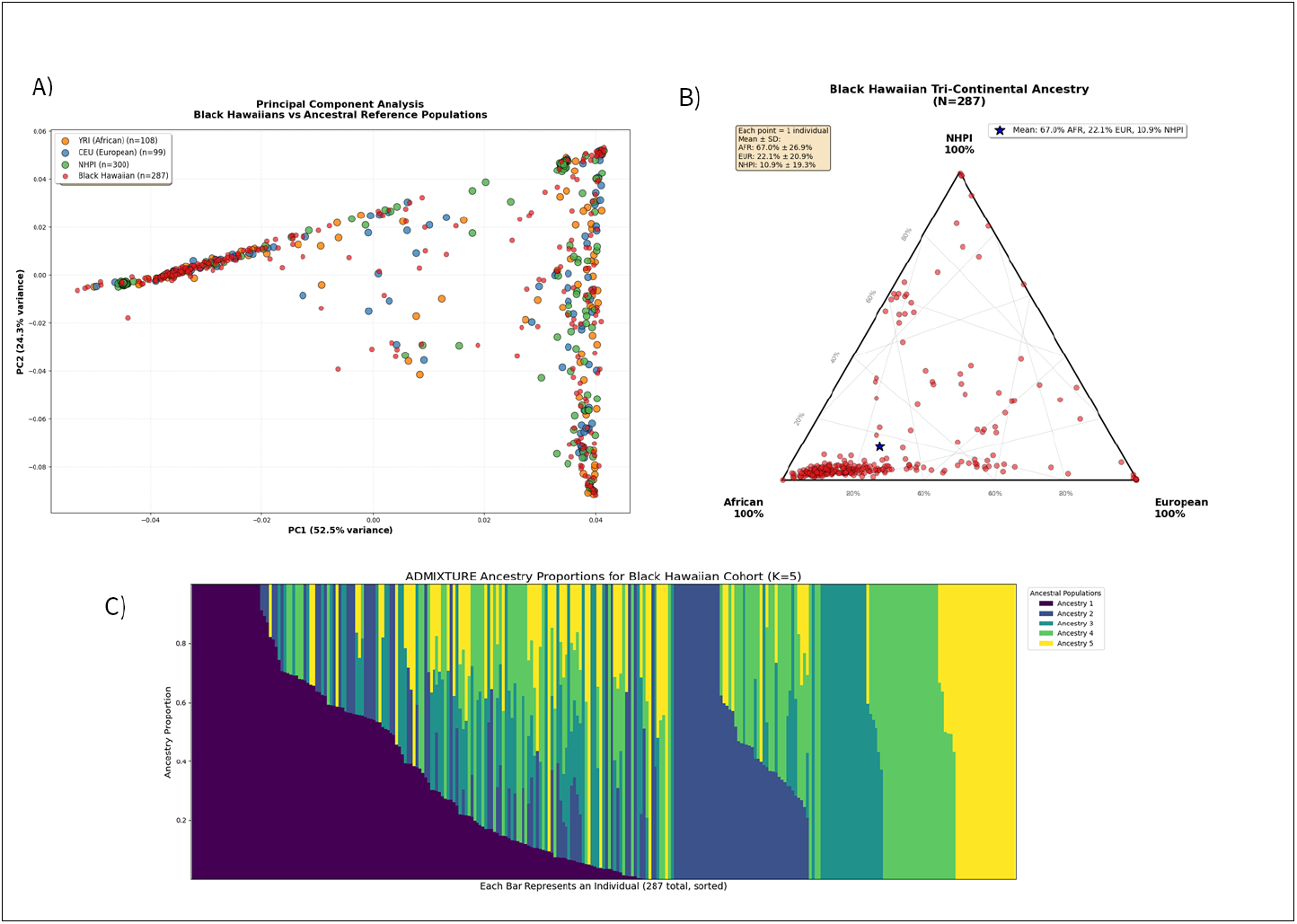
A) PCA plot showing Black Hawaiians relative to reference populations. B) Ternary plot of African-European-NHPI ancestry from RFMix. C) ADMIXTURE bar plot (K=5) showing individual ancestry proportions.

**Figure 3.**
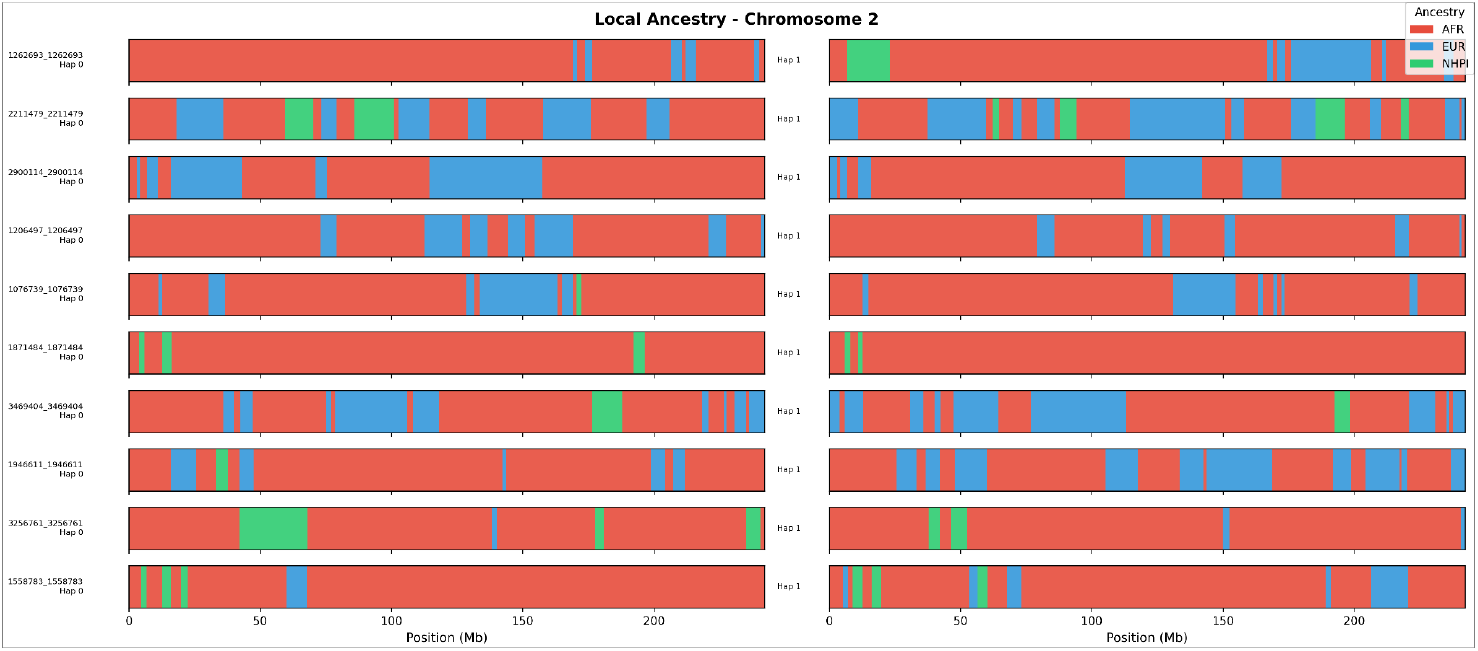
Local ancestry inference on chromosome 2 for representative Black Hawaiian individuals. The visualization displays phased haplotypes (“chromosome painting”), where colored segments represent distinct ancestral origins estimated by RFMix. The mosaic pattern illustrates the complex intercalation of African (red), European (blue), and Native Hawaiian/Pacific Islander (green) ancestry tracts.

### 3.3. Recent Effective Population Size Contraction

Demographic modeling using SMC++ revealed a distinct historical trajectory for the BH population characterized by a significant contraction in effective population size *N*_*e*_ in recent generations (Figure 4). The model estimated a current effective population size *N*_*e*_ of approximately 4,000 individuals. The demographic reconstruction shows a period of relative stability (*N*_*e*_ ≃6,000 - 7,000) extending from 600 years ago, back to approximately 100,000 years ago. This was preceded by a significant ancestral expansion event (peaking at *N*_*e*_ ≃20,000) roughly 150,000–300,000 years ago, consistent with the “Out of Africa” expansion signatures often observed in populations with substantial African ancestry. Most notably, the trajectory exhibits a sharp bottleneck within the last 10–20 generations (approx. 300–600 years), where *N*_*e*_ dropped from its stable plateau of ∼6,000 to the current ∼4,000. This recent decline likely captures the founder effects and Native Hawaiian depopulation following western contact and the specific demographic history of the BH population’s formation^27^.

**Figure 4.**
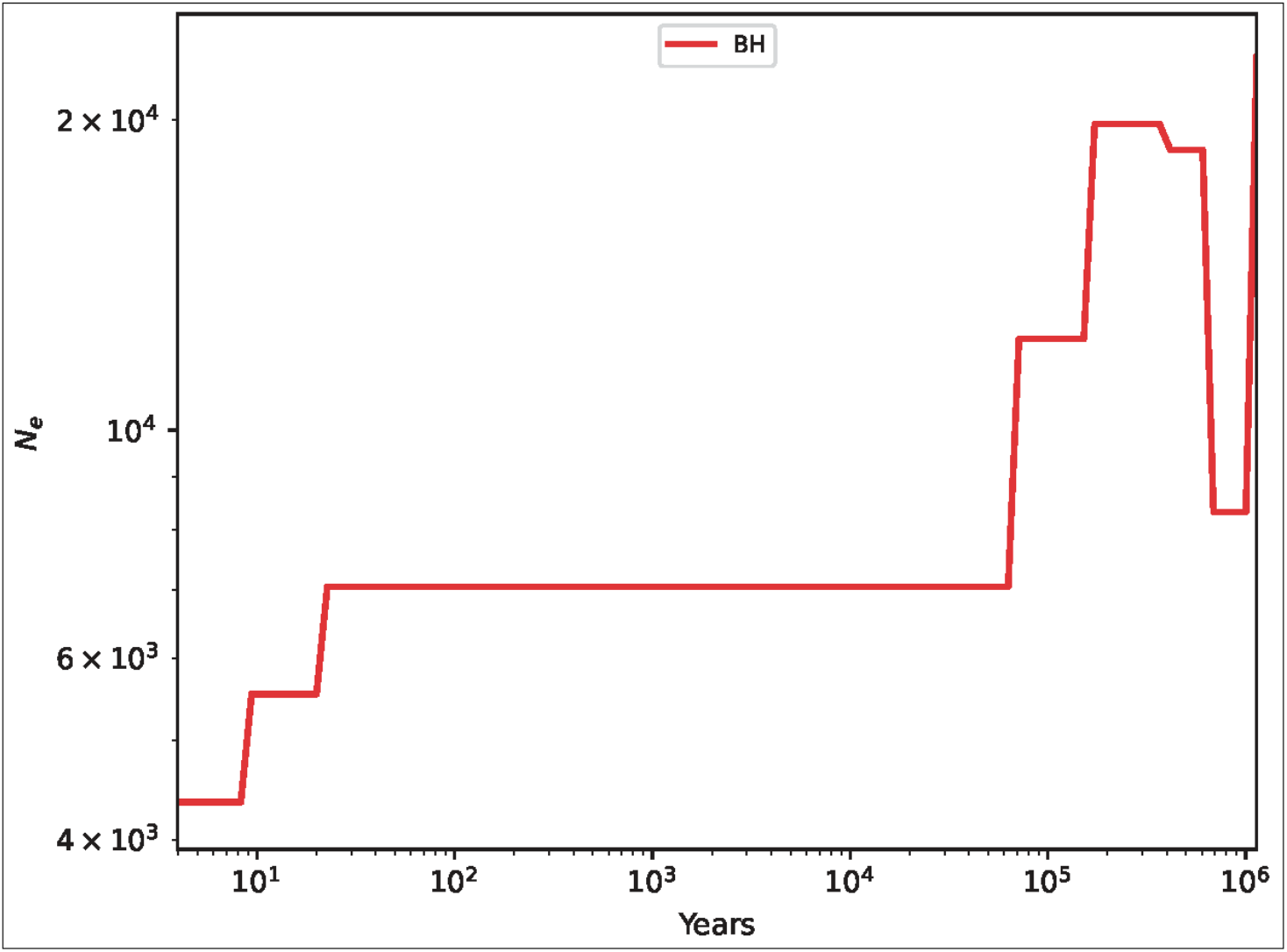
Demographic History of the Black Hawaiian Population. Step-plot showing the effective population size *N*_*e*_, y-axis, log scale) over time (years ago, x-axis, log scale, origin represents the current time) inferred by SMC++. The red line represents the Black Hawaiian (BH) cohort. The trajectory illustrates a large ancestral population size followed by a recent, sharp decline in *N*_*e*_ (bottleneck) closer to the present day 10–100 years ago).

### 3.4. Systematic Allele Frequency Differentiation

Genome-wide allele frequency comparison identified 8,909,583 variants present in BH genomes and all three reference populations. Mean absolute allele frequency differences were Δ=0.041 versus African (YRI), Δ=0.069 versus NHPI, and Δ=0.084 versus European (CEU), indicating that BH are genetically most similar to African populations but exhibit substantial differentiation from all three ancestral groups (Figure 5-A). The top 100 most differentiated variants (maximum Δ ranging from 0.555 to 0.613) were distributed across 15 chromosomes, with notable enrichment on chromosomes 3, 4, 10, 12, 15, and 17 (Figure 5-B, Table 2). Of these 100 variants, 42 (42%) mapped to 31 unique genes, while 58 (58%) were intergenic. The most frequently identified genes were *MYO9A* (4 variants), *RAB37* (4 variants), *LOC100131532* (4 variants), and *HERC2* (3 variants). Additional genes with multiple differentiated variants included *GPR158, CD300LF, EIF4B, DCUN1D1, PEAR1*, and *PLEKHA5*.

**Figure 5.**
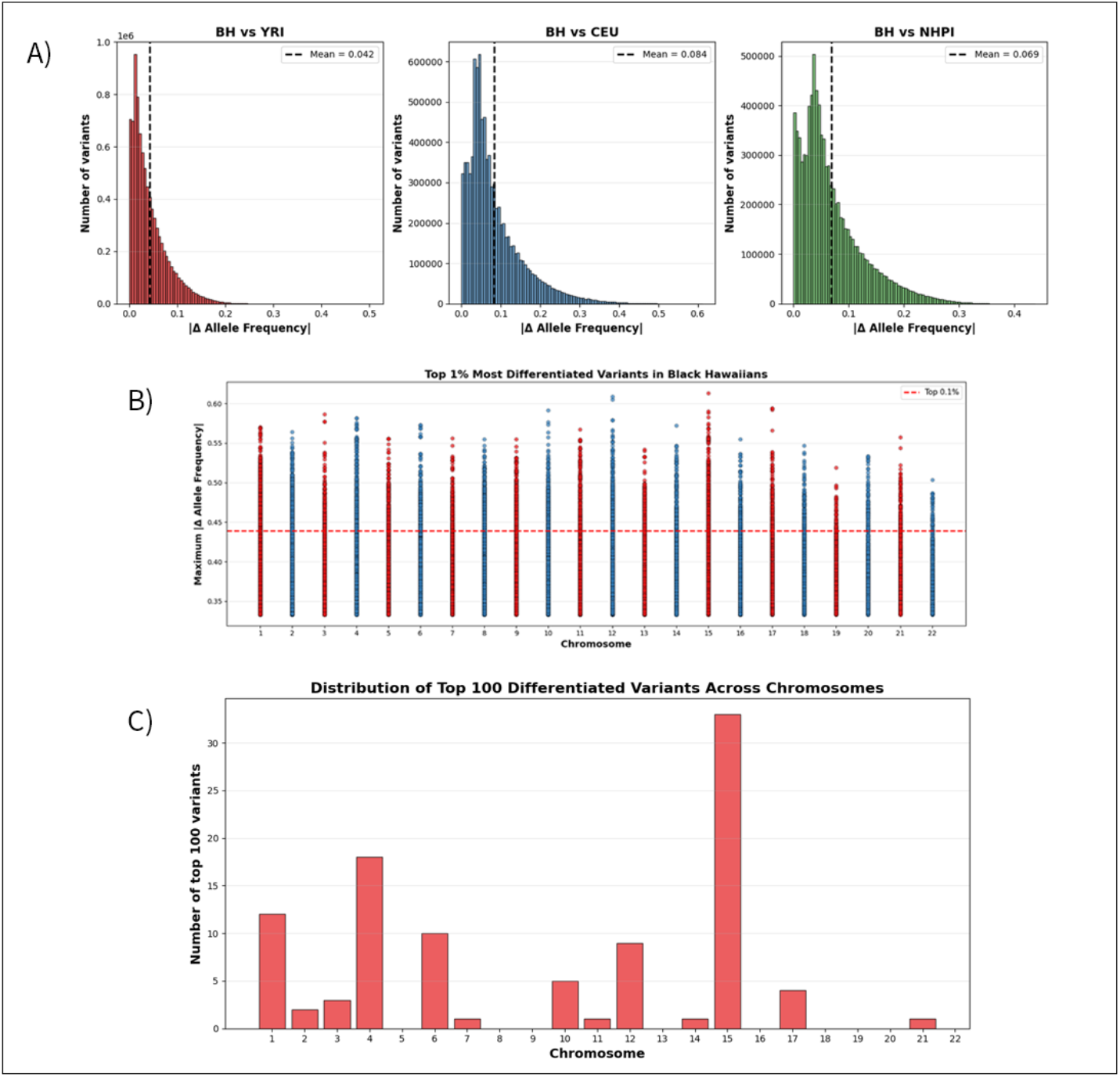
A) Distribution of allele frequency differences (Δ) for BH vs YRI, CEU, and NHPI. B) Manhattan plot showing top 1% most differentiated variants across chromosomes. C) Chromosome enrichment showing number of top 100 variants per chromosome.

### 3.5. Candidate Genes for Cardiometabolic Traits

To identify population-specific drivers of cardiometabolic risk, we prioritized variants within the top 1% of differentiation (Δ > 0.35) that map to genes with established roles in cardiovascular or metabolic physiology. We identified high-confidence candidate genes in two distinct physiological domains exhibiting extreme frequency divergence between BH and ancestral populations.

#### 3.5.1 Renal and Vascular Integrity (*MYO9A, PLEKHA5, PEAR1*)

The strongest signal for renal physiology was identified in *MYO9A* (Myosin IXA) on chromosome 15. We detected four variants in *MYO9A* within the top 100 most differentiated loci (Max Δ > 0.55). Biologically, *MYO9A* encodes a *Rho-GTPase* activating protein (Rho-GAP) critical for regulating cytoskeletal dynamics in podocytes^8,9^. Additionally, we identified highly differentiated variants in *PLEKHA5* (Pleckstrin Homology Domain Containing A5) and *PEAR1* (Platelet Endothelial Aggregation Receptor 1). *PLEKHA5* is implicated in the regulation of arterial stiffness via collagen cross-linking^10^, while *PEAR1* variants are associated with platelet hyper-reactivity and potential aspirin resistance^16^.

#### 3.5.2 Metabolic Regulation and Insulin Secretion (*RAB37, GPR158*)

In the metabolic domain, significant differentiation was observed in *RAB37* (Ras-related protein Rab-37) on chromosome 17, with four variants ranking in the top 100 most differentiated loci. *RAB37* is a key regulator of secretory autophagy in pancreatic β-cells, a pathway essential for high-volume insulin secretion under conditions of metabolic stress^12^. We also identified differentiation in *GPR158* (G Protein-Coupled Receptor 158). *GPR158* is linked to central energy homeostasis and osteocalcin signaling^14^, with the BH cohort exhibiting significantly higher frequencies of variants associated with energy conservation compared to African reference panels.

## 4. Discussion

This study represents the first comprehensive genomic characterization of BH individuals, revealing complex tri-continental admixture patterns, elevated cardiometabolic burden, and systematic allele frequency differentiation from ancestral populations. Our findings establish foundational genomic resources for this underrepresented population and demonstrate the critical importance of disaggregated research approaches in admixed communities.

### 4.1. Tri-Continental Admixture and Population Heterogeneity

BH exhibit substantial African ancestry (67%) alongside meaningful European (22%) and NHPI (11%) components, with high inter-individual variation reflecting diverse admixture histories. This ancestry pattern distinguishes BH from both African Americans (typically 75-85% African ancestry with minimal NHPI) and other NHPI groups (typically <5% African ancestry). The absence of discrete sub-clusters within our cohort suggests continuous admixture rather than distinct foundational populations, potentially reflecting ongoing gene flow and complex demographic history in Hawai’i. The high standard deviations in ancestry proportions (26.9% for African component) demonstrates that treating BH as a monolithic genetic entity would obscure individual disease risks. Therefore, future research must move beyond broad labels to utilize continuous ancestry metrics, ensuring that the diversity within this unique population (and similar case in other populations) is accurately reflected in precision medicine. This heterogeneity has important implications for genetic association studies, as individuals with varying ancestry proportions may have different genetic risk profiles. Local ancestry inference provides a framework for accounting for this variation in future association analyses by enabling ancestry-specific effect estimates.

### 4.2. Demographic History and Founder Effects

Beyond static admixture proportions, our SMC++ analysis provides the first temporal reconstruction of the population demographic history of BH. The demographic trajectory mirrors the known anthropological history of the region: a long period of stability (N_e_ ≃6,000) followed by a sharp, recent contraction in effective population size (N_e_ ≃4,000$) within the last 10–20 generations. This distinct “bottleneck” likely captures the coalescence of the BH population through the admixture of relatively small founding groups, European explorers, Asian and Pacific laborers, and African individuals, alongside the tragic historical depopulation of Native Hawaiians following

Western contact^19^. Crucially, this recent bottleneck provides the population-genetic context for the high allele frequency differentiation we observed. In populations with reduced effective size (*N*_*e*_), genetic drift exerts a stronger influence than in large, panmictic populations. This suggests that the high frequency of risk alleles in *MYO9A* and *RAB37* may have been driven to prominence by founder effects, where rare variants carried by a few ancestors rose in frequency by chance during the population’s formation, rather than solely by adaptive selection. This “Founder Effect” architecture is characteristic of island populations and underscores the necessity of population-specific reference panels, as variants rare in the continental diaspora may be common and clinically relevant in Hawai’i.

### 4.3. Elevated Cardiometabolic Burden and Health Disparities

The prevalence of hypertension (34.8%) and obesity (mean BMI 31.2) in our BH cohort exceeds national averages and parallels rates observed in both African American and Native Hawaiian populations. This convergence of risk likely reflects a complex interplay between socioeconomic factors and the unique genetic architecture identified in this study. While the relatively lower prevalence of T2D (7.7%) and cardiovascular disease (6.3%) may reflect the younger age distribution of the AoU cohort or underdiagnosis, the high frequency of risk alleles in metabolic regulators like *RAB37* and *GPR158* suggests a substantial latent genetic risk that may manifest with age or environmental stress. These findings emphasize that standard clinical risk models may be insufficient for this population. Crucially, our identification of distinct candidate loci challenges the reliance on ancestral surrogates; simply screening for African-specific markers (e.g., *APOL1*) or European Polygenic Risk Scores (PRS) would miss the population-specific drivers, such as *MYO9A*-mediated cytoskeletal defects, that appear central to the BH health burden.

### 4.4. Allele Frequency Differentiation and Candidate Genes

The identification of highly differentiated variants in genes with specific mechanistic roles suggests that the high cardiometabolic burden in BH is not merely a reflection of generalized admixture, but potentially driven by distinct, population-specific biological pathways (Figure 6).

**Figure 6.**
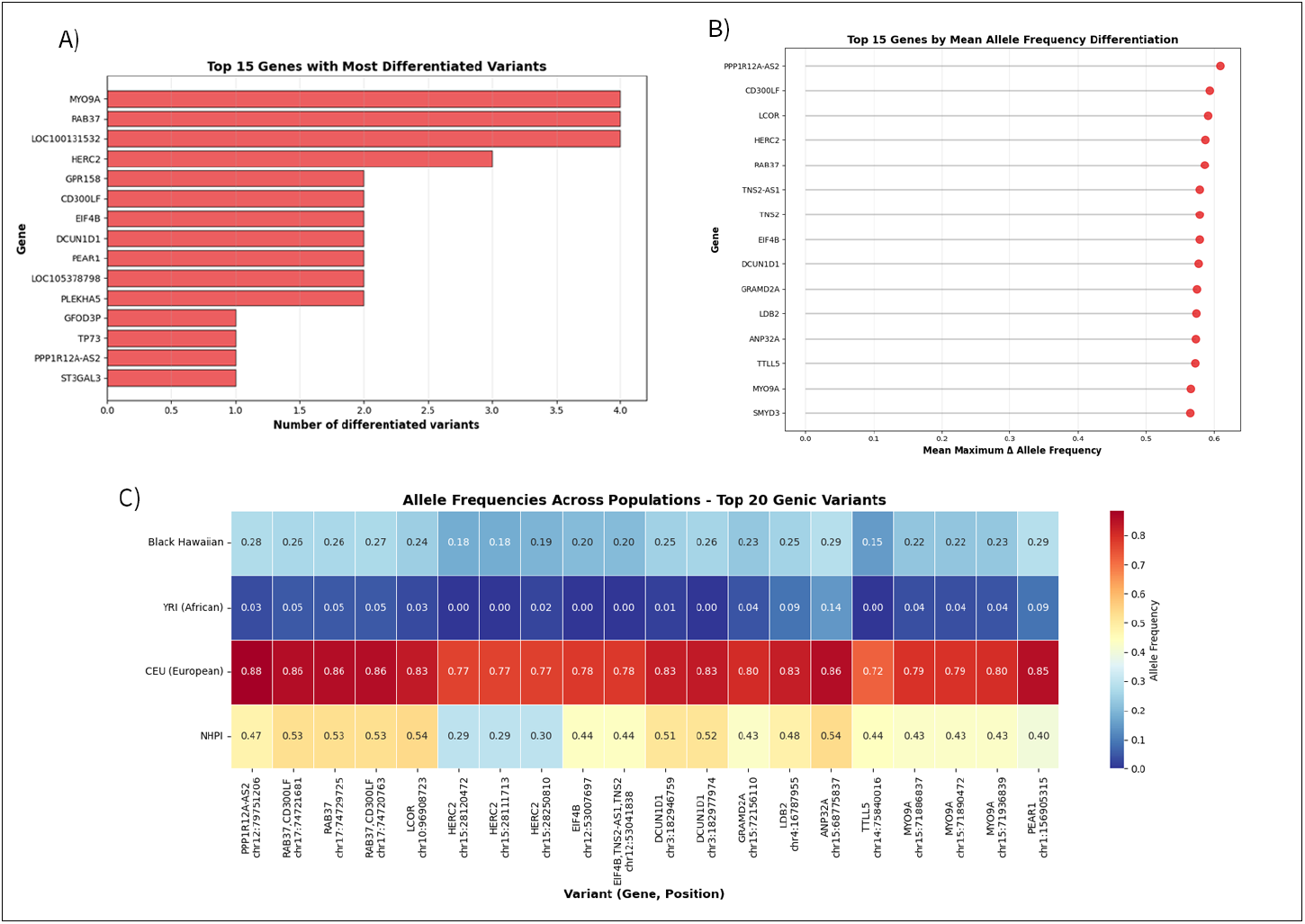
A) Top 15 genes ranked by number of differentiated variants. B) Mean differentiation score for top genes. C) Heatmap showing allele frequencies across populations for top 20 genic variants.

#### 4.4.1. A Mechanostructural Hypothesis for Renal Disease

Our findings challenge the prevailing paradigm of kidney disease in African-admixed populations. While *APOL1* cytotoxicity is the dominant risk factor in African Americans, the identification of *MYO9A* as the top candidate in BH suggests a distinct etiology^8,9^. *MYO9A* functions as a Rho-GAP essential for podocyte cytoskeletal plasticity; the variants identified here likely impair the “braking” of RhoA, leading to constitutive actin stress fiber formation and cytoskeletal rigidity [1,3]. In the context of the high hypertension rates observed in this cohort (34.8%), this creates a “Mechanostructural Vulnerability”: rigid podocytes unable to flex under hemodynamic stress, leading to physical effacement rather than chemical toxicity^10^. This structural hypothesis is further supported by the differentiation of *PLEKHA5*. Mechanistically, *PLEKHA5* recruits *PDZD11* to traffic the copper transporter *ATP7A* to the cell surface, a step required to activate Lysyl Oxidase (LOX) for elastin cross-linking^11,12^. Dysregulation of this complex suggests that hypertension in this population may be driven by vascular stiffness and oxidative remodeling^13^, a pathology distinct from the salt-sensitive, volume-dependent hypertension typically managed with diuretics in other populations.

#### 4.4.2. The Two-Hit Metabolic Model

For T2D, our data supports a sophisticated “Two-Hit” model of disease progression. (1) Susceptibility (The First Hit): The high frequency of *GPR158* variants suggests a genetic predisposition toward efficient energy storage. Mechanistically, *GPR158* functions as the central receptor for osteocalcin, a bone-derived hormone that regulates appetite, stress responses, and metabolic homeostasis in the brain^14^. Dysregulation of this “Bone-Brain-Metabolic Axis” aligns with the high prevalence of obesity (mean BMI 31.2) and suggests that skeletal physiology may be driving metabolic outcomes in this population. (2) Failure (The Second Hit): The differentiation of *RAB37* provides the mechanism of decompensation. *RAB37* acts as the master regulator of secretory autophagy, a specialized “emergency” pathway required for high-volume insulin secretion under metabolic stress^15^. We hypothesize that the identified variants impair this critical response, rendering pancreatic β-cells unable to meet the heightened insulin demand driven by obesity, leading to rapid β-cell exhaustion and overt diabetes.

#### 4.4.3 Pharmacogenomic Implications

Finally, the differentiation of *PEAR1* has immediate clinical relevance. Biologically, *PEAR1* is a transmembrane receptor responsible for stabilizing platelet aggregates upon cell-cell contact^16^. The variants identified in this cohort are strongly linked to platelet hyper-reactivity and “Aspirin Resistance”, a phenomenon where standard COX-1 inhibition fails to suppress thrombus formation because the *PEAR1* signaling pathway remains active^16,17^. Given that low-dose aspirin is the standard of care for cardiovascular prevention, a significant segment of the BH population may be receiving suboptimal prophylaxis. This highlights a critical opportunity for precision medicine: the potential necessity of investigation into *P2Y12* inhibitors (e.g., clopidogrel) as an alternative therapy, particularly given that *PEAR1* functions as a specific modulator of platelet signaling rather than a general cardiovascular susceptibility gene^17,18,19^.

#### 4.5. Implications for Equitable Precision Medicine

Our findings underscore fundamental challenges in applying genetic risk prediction models developed in single-ancestry populations to admixed individuals. Standard PRS derived exclusively from European or West African cohorts likely suffer from reduced accuracy in BH due to the unique allele frequency distributions (mean Δ=0.041-0.084) and distinct LD patterns created by tri-continental admixture. The local ancestry inference framework established here provides a blueprint for overcoming these limitations. Through the explicit modeling of the mosaic of African, European, and Pacific ancestry, future association studies can leverage admixture mapping to detect causal variants that would otherwise remain hidden in aggregated datasets. As sample sizes increase through continued recruitment of diverse populations into biobanks like All of Us, such disaggregated approaches will be essential to ensure that the promises of precision medicine, from accurate risk prediction to pharmacogenomic tailoring, are equitably realized for complexly admixed communities.

### 5. Limitations

Several limitations warrant consideration.

1. Our sample size (n=287), while representing the first analysis of BH, is modest for genetic association studies and precludes detection of variants with small effect sizes.
2. Reliance on self-identified NHPI individuals from *All of Us* as a reference panel introduces potential ‘proxy bias.’ Given the extensive post-contact admixture history in Hawai’i, these reference individuals likely harbor cryptic European or Asian ancestry tracts. Consequently, our local ancestry inference (RFMix) may have underestimated European admixture in the BH cohort, as European haplotypes could be misclassified as ‘NHPI-like’ if they match the admixed reference panel. While we mitigated this by using a consensus of 300 individuals, future studies should prioritize the use of ancient DNA or highly curated indigenous reference panels (e.g., from the Consortia for Polynesian Genomics) to improve the resolution of Pacific ancestry tracts.
3. Phenotype data were extracted from electronic health records and may be incomplete or subject to ascertainment bias.
4. Self-reported race and ethnicity categories, while socially meaningful, represent imperfect proxies for genetic ancestry. Some individuals identifying as BH may have limited NHPI genetic ancestry, while others with substantial NHPI ancestry may not self-identify using this label. Future research should investigate the concordance between self-identification and genetic ancestry, recognizing that both are important dimensions of individual and community identity.
5. Regarding our allele frequency differentiation analysis, we interpreted high differentiation as a signal of potential population-specific importance. However, we acknowledge that deviation from ancestral allele frequencies can result from two distinct processes: (1) admixture dynamics, where the observed frequency simply reflects the weighted average of ancestral proportions, and (2) post-admixture evolutionary forces such as genetic drift or selection. While we did not establish a simulated ‘baseline admixture’ null model to disentangle these effects, the magnitude of differentiation observed for top candidates like *MYO9A* (Δ > 0.5) exceeds what would typically be expected from simple linear admixture, particularly given the recent population bottleneck we identified. Future studies utilizing local ancestry-aware simulation frameworks will be necessary to definitively distinguish between admixture-induced drift and positive selection.
6. To maximize the available sample size of this underrepresented population (N=287), strict kinship filtering was not applied prior to population structure analyses. While this approach preserves statistical power for allele frequency differentiation, it allows for the potential retention of cryptic relatedness, which could subtly influence fine-scale clustering patterns or effective population size estimates. Future association studies in this cohort will require linear mixed models or kinship pruning to control for population stratification.

### 6. Future Directions

This foundational study establishes several priorities for future research in BH genomics:

1. **Functional Validation of Candidate Mechanisms:** Future experimental work must move beyond association to causation. Specifically, this includes assessing podocyte cytoskeletal rigidity in *MYO9A*-variant cell lines under flow stress and quantifying secretory autophagy efficiency in *RAB37*-edited β-cells to definitively link these variants to organ failure.
2. **Admixture Mapping and Fine-Mapping:** Leveraging the unique tri-continental local ancestry architecture established here (Figure 2) to fine-map disease loci that are invisible in homogeneous populations. This approach is particularly powerful for traits where risk alleles are ancestry-specific, such as the *PEAR1* pharmacogenomic signal.
3. **Ancestry-Aware Polygenic Risk Scores (PRS):** Developing and validating PRS models that explicitly incorporate local ancestry weighting. Standard “pan-African” or “European” scores likely fail in this population; a BH-specific PRS that accounts for the mosaic genome is essential for clinical utility.
4. **Gene-Environment and Social Interaction:** Investigating how the identified genetic drivers (e.g., the *GPR158-Osteocalcin* axis) interact with social determinants of health. Understanding whether metabolic genetic risk is amplified by specific environmental stressors will be key to designing holistic interventions.

## Conclusions

This study provides the first comprehensive genomic characterization of Black Hawaiian individuals, revealing complex admixture patterns, substantial cardiometabolic burden, and systematic genetic differentiation from ancestral populations. Beyond description, our identification of distinct mechanistic drivers, including *MYO9A* (renal cytoskeletal integrity), *RAB37* (secretory autophagy), and *PEAR1* (platelet signaling), challenges current biomedical paradigms and provides concrete targets for future functional validation.

Critically, our findings demonstrate that Black Hawaiians occupy a distinct region of genetic space that cannot be adequately represented by aggregation with African American or Pacific Islander cohorts. As precision medicine initiatives expand, ensuring equitable inclusion of such admixed populations is essential for reducing health disparities. The genomic resources and analytical frameworks established here provide a model for characterizing other underrepresented populations, advancing the goal of a genomic medicine that is not only precise but truly inclusive.

## Acknowledgments

This research was conducted using data from the *All of Us* Research Program, supported by the National Institutes of Health (NIH). We thank the *All of Us* participants for their contributions. We acknowledge the 1000 Genomes Project for providing reference panel data.

## Data Availability

Individual-level genotype and phenotype data are available through the NIH *All of Us* Research Program (https://www.researchallofus.org/) subject to appropriate data use agreements. The code used for this analysis is available to authorized users on the *All of Us* Researcher Workbench in the workspace titled “BH STUDY”. Readers can register for access at ResearchAllofUs.org.

## References

1. Danecek P, Bonfield JK, Liddle J, et al. Twelve years of SAMtools and BCFtools. Gigascience. 2021;10(2):giab008. doi:10.1093/gigascience/giab008

2. Purcell S, Neale B, Todd-Brown K, et al. PLINK: a tool set for whole-genome association and population-based linkage analyses. Am J Hum Genet. 2007;81(3):559–575. doi:10.1086/519795

3. Hofmeister RJ, Ribeiro DM, Rubinacci S, Delaneau O. Accurate rare variant phasing of whole-genome and whole-exome sequencing data in the UK Biobank. Nat Genet. 2023;55(7):1243–1249. doi:10.1038/s41588-023-01415-w

4. The 1000 Genomes Project Consortium. A global reference for human genetic variation. Nature. 2015;526(7571):68–74. doi:10.1038/nature15393

5. Alexander DH, Novembre J, Lange K. Fast model-based estimation of ancestry in unrelated individuals. Genome Res. 2009;19(9):1655–1664. doi:10.1101/gr.094052.109

6. Maples BK, Gravel S, Kenny EE, Bustamante CD. RFMix: a discriminative modeling approach for rapid and robust local-ancestry inference. Am J Hum Genet. 2013;93(2):278–288. doi:10.1016/j.ajhg.2013.06.020

7. Terhorst, J., Kamm, J. & Song, Y. Robust and scalable inference of population history from hundreds of unphased whole genomes. Nat Genet 49, 303–309 (2017). 10.1038/ng.3748

8. Li Q, Gulati A, Lemaire M, et al. Rho-GTPase activating protein myosin MYO9A identified as a novel candidate gene for monogenic focal segmental glomerulosclerosis. Kidney Int. 2021;99(5):1102–1117. doi:10.1016/j.kint.2020.12.022

9. Schlondorff JS. My, oh, MYO9A! Just how complex can regulation of the podocyte actin cytoskeleton get? Kidney Int. 2021;99(5):1065–1067. doi:10.1016/j.kint.2021.02.006

10. Zhu L, Jiang R, Aoudjit L, Jones N, Takano T. Activation of RhoA in podocytes induces focal segmental glomerulosclerosis. J Am Soc Nephrol. 2011;22(9):1621–1630. doi:10.1681/ASN.2010111146

11. Sluysmans S, Méan I, Xiao T, et al. PLEKHA5, PLEKHA6, and PLEKHA7 bind to PDZD11 to target the Menkes ATPase ATP7A to the cell periphery and regulate copper homeostasis. Mol Biol Cell. 2021;32(21):ar34. doi:10.1091/mbc.E21-07-0355

12. Martínez-González J, Varona S, Cañes L, et al. Emerging roles of lysyl oxidases in the cardiovascular system: new concepts and therapeutic challenges. Biomolecules. 2019;9(10):610. doi:10.3390/biom9100610

13. Martínez-Revelles S, García-Redondo AB, Avendaño MS, et al. Lysyl oxidase induces vascular oxidative stress and contributes to arterial stiffness and abnormal elastin structure in hypertension: role of p38MAPK. Antioxid Redox Signal. 2017;27(6):379–397. doi:10.1089/ars.2016.6642

14. Khrimian L, Obri A, Ramos-Brossier M, et al. Gpr158 mediates osteocalcin’s regulation of cognition. J Exp Med. 2017;214(10):2859–2873. doi:10.1084/jem.20171320

15. Wu SY, Wu HT, Wang YC, et al. Secretory autophagy promotes RAB37-mediated insulin secretion under glucose stimulation both in vitro and in vivo. Autophagy. 2023;19(4):1239–1257. doi:10.1080/15548627.2022.2123098

16. Lewis JP, Ryan K, O’Connell JR, et al. Genetic variation in PEAR1 is associated with platelet aggregation and cardiovascular outcomes. Circ Cardiovasc Genet. 2013;6(2):184–192. doi:10.1161/CIRCGENETICS.111.964627

17. Khan H, Kanny O, Syed MH, Qadura M. Aspirin resistance in vascular disease: a review highlighting the critical need for improved point-of-care testing and personalized therapy. Int J Mol Sci. 2022;23(19):11317. doi:10.3390/ijms231911317

18. Lewis JP, Riaz M, Xie S, et al. Genetic variation in PEAR1, cardiovascular outcomes and effects of aspirin in a healthy elderly population. Clin Pharmacol Ther. 2020;108(6):1289–1298. doi:10.1002/cpt.1959

19. Yang WY, Petit T, Cauwenberghs N, et al. PEAR1 is not a major susceptibility gene for cardiovascular disease in a Flemish population. BMC Med Genet. 2017;18(1):45. doi:10.1186/s12881-017-0411-x

20. Schmitt RC. Demographic Statistics of Hawaii: 1778–1965. Honolulu (HI): University of Hawaii Press; 1968.

21. Xu K, Ye S, Zhang S, et al. Impact of Platelet Endothelial Aggregation Receptor-1 genotypes on platelet reactivity and early cardiovascular outcomes in patients undergoing percutaneous coronary intervention and treated with aspirin and clopidogrel. Circ Cardiovasc Interv. 2019;12(5):e007019. doi:10.1161/CIRCINTERVENTIONS.118.007019

22. Sun H, Lin M, Russell EM, et al. The impact of global and local Polynesian genetic ancestry on complex traits in Native Hawaiians. PLoS Genet. 2021;17(2):e1009273. doi:10.1371/journal.pgen.1009273

23. Chiang CWK, Marcus JH, Sidore C, et al. The impact of global and local Polynesian genetic ancestry on complex traits in Native Hawaiians. PLoS Genet. 2021;17(2):e1009273.

